# Impact of central complex lesions on visual orientation in ants: Turning behaviour, but not the overall movement direction, is disrupted

**DOI:** 10.1101/2021.02.02.429334

**Authors:** Scarlett Dell-Cronin, Cornelia Buehlmann, Angela Diyalagoda Pathirannahelage, Roman Goulard, Barbara Webb, Jeremy Niven, Paul Graham

## Abstract

Wood ants are excellent navigators using a combination of innate and learnt navigational strategies to travel between their nest and feeding sites. Visual navigation in ants has been studied extensively, however, we only know little about the underlying neural mechanisms. The central complex (CX) is located at the midline of the insect brain. It receives sensory input that allows an insect to keep track of the direction of sensory cues relative to its own orientation and to control movement. We show here direct evidence for the involvement of the central complex in the innate visual orientation response of freely moving wood ants. Lesions in the CX disrupted the control of turning in a lateralised manner, but had no effect on the overall heading direction, walking speed or path straightness.

## INTRODUCTION

Ant foragers travel diligently back and forth between their nest and feeding sites using a combination of innate and learnt navigational strategies (Knaden & Graham 2016; Buehlmann *et al.* 2020b). Innate strategies such as path integration (e.g. (Collett *et al.* 2001; Mueller & Wehner 2010)), pheromone trails (e.g. (Harrison *et al.* 1989)), attraction to food odours (Buehlmann *et al.* 2014), and/or attraction to visual cues (e.g. (Graham *et al.* 2003; Collett 2010)) are vital for ants that are unfamiliar in an environment. Wood ants show an innate attraction to large conspicuous objects which can form part of their navigational repertoire (Voss 1967; Graham *et al.* 2003; Buehlmann *et al.* 2020a; Buehlmann *et al.* 2020c; Buehlmann & Graham 2021). This innate behaviour can form the basis for learnt foraging routes (Graham *et al.* 2003). Visual navigation in ants has been studied extensively (Zeil 2012; Collett *et al.* 2013; Wehner *et al.* 2014; Graham & Philippides 2017), however, knowledge about the underlying neural mechanisms lags behind (Webb & Wystrach 2016; Heinze 2017).

The central complex (CX), a collection of neuropils located at the midline of the insect brain (Strausfeld 1999), receives sensory input allowing an insect to keep track of the direction of sensory cues relative to its own orientation (Seelig & Jayaraman 2015), and to directly control movement (Martin *et al.* 2015). A few studies have shown that the CX plays a role in insects that navigate using celestial information (e.g. locusts: (Vitzthum *et al.* 2002; Pegel *et al.* 2018); dung beetles: (el Jundi *et al.* 2015); monarch butterfly: (Heinze & Reppert 2011; Heinze *et al.* 2013); bees: (Stone *et al.* 2017); crickets: (Sakura *et al.* 2008)) or terrestrial visual information (fruit flies: (Liu *et al.* 2006; Neuser *et al.* 2008; Ofstad *et al.* 2011; Kuntz *et al.* 2012; Seelig & Jayaraman 2013); cockroaches: (Ritzmann *et al.* 2008); locust: (Rosner & Homberg 2013)). In these insects, the CX has a role in the control of speed, turning behaviour, and spatial orientation (reviewed in (Pfeiffer & Homberg 2014; Varga *et al.* 2017; Honkanen *et al.* 2019)). Furthermore, an increase of the CX volume was shown in new ant foragers that were exposed to skylight information during the learning of navigationally relevant information at the offset of their foraging career, indicating its involvement in navigation (Grob *et al.* 2017).

We show here direct evidence for the involvement of the CX in the innate visual orientation response of freely moving wood ants. Lesions in the CX disrupted the control of turning in a lateralised manner, but had no effect on the overall heading direction, walking speed or path straightness.

## METHODS

### Ants

Experiments were performed with laboratory kept wood ants *Formica rufa L.* collected from Ashdown forest, East Sussex, UK. Ants were kept in the laboratory under a 12 h light: 12 h darkness cycle at 25-27° C. Ants were fed *ad libitum* with sucrose, dead crickets and water. During the experiments, food was limited to a minimum to increase the ants’ foraging motivation, but water was permanently available.

### Lesion procedures

Ants were immobilised on ice for 90 seconds and harnessed in a custom-made holder keeping their head fixed with plasticine while their body was free to move. Antennae were restrained with a pin. To access the CX, a small window was cut with a piece of razor blade below the medial ocellus. Mechanical lesions were made relative to landmarks (including tracheal branches) visible on the anterior surface of the brain. The bottom of the central tracheal branch (Figure 1A), directly below the medial ocellus, indicated the position of the CX (Figure 1B). Glass capillaries (Harvard Apparatus, Cambourne, UK; 30-0035; 1.00 mm outer diameter, 0.78 mm inner diameter, 10 mm length) were pulled with a P-97 Micropipette Puller (Sutter Instrument, Novato, California, USA) and then broken manually to a tip size of 10 µm and dipped in black ink to aid visibility. After inserting the capillary into either the left or right side of the CX (Figure 1A), the cuticle lid was replaced and secured in place with a small drop of cyanoacrylate adhesive. Control ants were handled and dissected in the same way as lesion ants except for the actual lesion.

**Figure 1:**
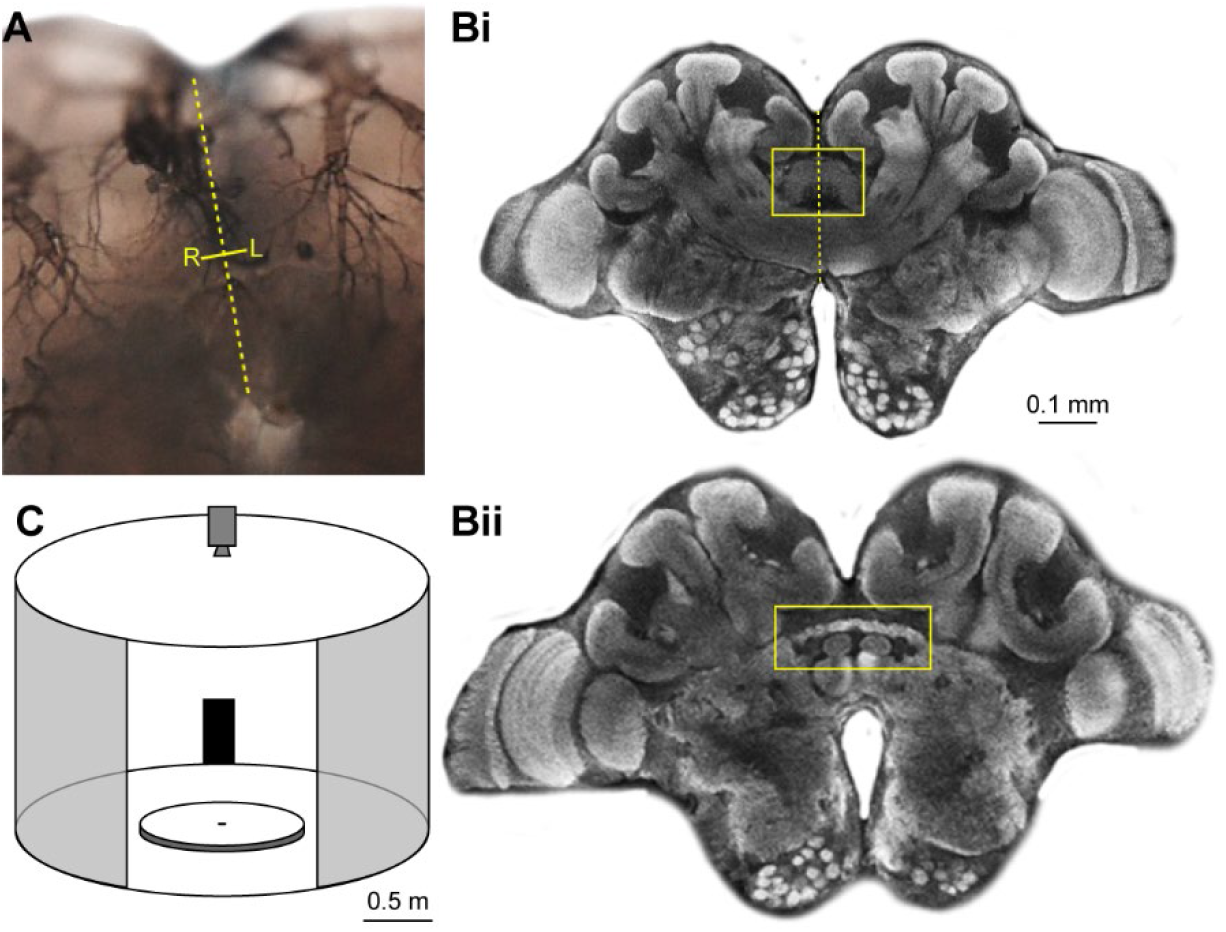
CX lesion locations and experimental setup. **(A)** Mechanical lesions were made relative to landmarks (including tracheal branches) visible on the anterior surface of the brain. The bottom of the central tracheal branch, directly below the medial ocellus, indicated the position of the CX. Yellow dashed line indicates brain midline and small yellow line with L (lesion on left) and R (lesion on right) show lesion locations. **(B)** Confocal scan of an anti-synapsin labelled wood ant brain. CX is marked with a yellow rectangle. **(C)** The experimental arena in which naïve ants were recorded. Circular white platform (radius: 60 cm) is located in the centre of a cylinder (radius: 1.5 m, height: 1.8 m). A 20° wide black rectangle (height: 90 cm, width: 52 cm) is mounted at the inner wall of the surrounding cylinder. A camera recorded the ants’ paths from above. A small door permitted access to the arena shown here open and larger for clarity.

### Experimental setup

After resting for 10 minutes, ants were released in the centre of a circular platform (120 cm in diameter) within a cylindrical arena (diameter 3 m, height 1.8 m) with white walls (Figure 1C). A 20° wide rectangle (height: 90 cm, width: 52 cm) was placed on the inner wall of the surrounding cylinder. To remove possible olfactory cues, the surface of the platform was covered with white paper, which was rotated between recordings. The centre of the platform consisted of a cylindrical release chamber of 6.5 cm diameter, which was remotely lowered to release the ant onto the platform. The ants’ position and body orientation were recorded every 20 ms using a tracking video camera (Trackit, SciTrackS GmbH) and trajectories were analysed in Matlab.

## RESULTS

### Walking speed and overall path straightness are not affected by CX lesions

CX lesions affected the ants’ propensity to move. 12 out of 80 CX-lesioned ants did not leave the release chamber (r = 3.25 cm), whereas only 1 out of 55 control ants, which had all the handling including head opening but no lesion, did not reach r = 3.25 cm (Chi-square test, p < 0.05). Walking speed and general path straightness of those ants that left the release chamber were not affected by the lesions (Control, n = 54 ants; CX lesions, n = 68 ants; Mann Whitney test; walking speed, p > 0.05; path straightness, p > 0.05). Hence, although initial tendency to move was lower in lesioned ants, CX lesions had no impact on walking speed or overall path straightness in ants that left the release chamber. However, even though there was no difference in overall path straightness, lesioned ants produced a significantly higher number of loops, i.e. path segments where an ant turned around and re-crossed its own path, than the control ants (Control, median, 4 loops/m; Lesions, median, 9 loops/m; Mann Whitney test, p < 0.05).

### Innate visual orientation of CX lesioned ants is not impaired

CX lesions did not affect the ants’ innate preference for large dark objects. Control ants (Figure 2A, n = 54 ants) and lesioned ants from both groups (Figure 2B, left lesions, LLes, n = 35 ants; Figure 2C, right lesion, RLes, n = 33 ants) were directed (Rayleigh test, all p < 0.001). Ants from the three groups did not differ from each other (Watson Williams tests, all p > 0.05) and they all approached the visual cue, i.e. the 95% CI of the ants’ final heading direction encompassed the visual cue (Figure 2).

**Figure 2:**
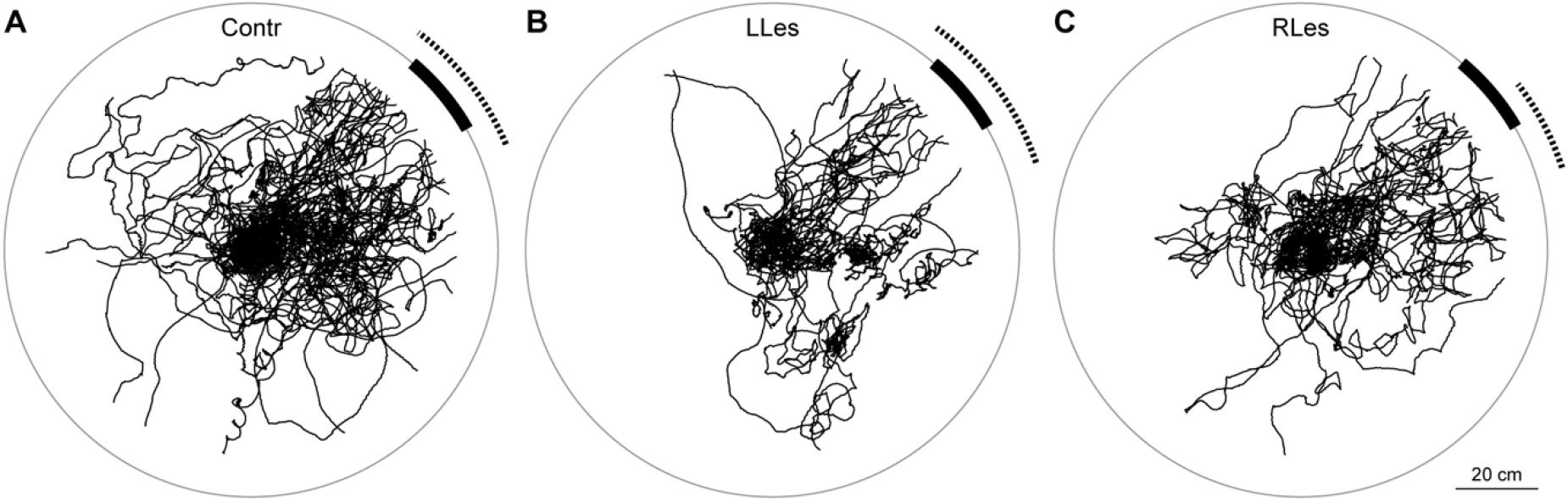
Innate visual attraction in wood ants is not affected by lesions in the CX. **(A)** Paths of control ants released at the centre of the arena are shown as black lines. Dotted arcs show 95% confidence intervals (CIs) of the heading directions. The visual cue is shown at the platform edge instead of on the cylinder wall. **(B)** As for A but for ants with lesions on the left side of the CX. **(C)** As in A but for ants with lesions on the right side of the CX. For sample sizes see Results.

### CX lesions disrupt turning

Detailed path analysis from ants that approached the centre of the visual cue +/− 90° revealed that control ants as expected spent on average an equal amount of time turning to the left and right (Contr, n = 42 ants; Wilcoxon test, p > 0.05), whereas the average length of left and right turns in both lesion groups was significantly different (LLes, n = 29 ants; RLes, n = 27 ants; Wilcoxon test, both p < 0.001). Lesioned ants spent less time turning to the ipsilateral side of the lesion, i.e. ants lesioned on the right spent less time turning to the right and vice versa (Figure 3A). Analysis of the magnitude of these turns revealed that there was no difference between left and right turns in all three groups (Figure 3B; Wilcoxon tests; Contr, n = 42 ants; LLes, n = 28 ants; RLes, n = 27 ants; all p > 0.05). The angles at the end of the turns differed in all three groups between left and right turns (Figure 3C). At the end of a right turn, ants looked to the right relative to the centre of the visual cue, whereas at the end of a left turn, ants looked to the left of the centre of the visual cue (Wilcoxon tests; Contr, n = 42 ants, p < 0.001; LLes, n = 28 ants, p < 0.01; RLes, n = 27 ants, p < 0.001). The end points after both the left and right turns respectively did not differ between the three groups (Watson Williams tests, all p > 0.05).

**Figure 3:**
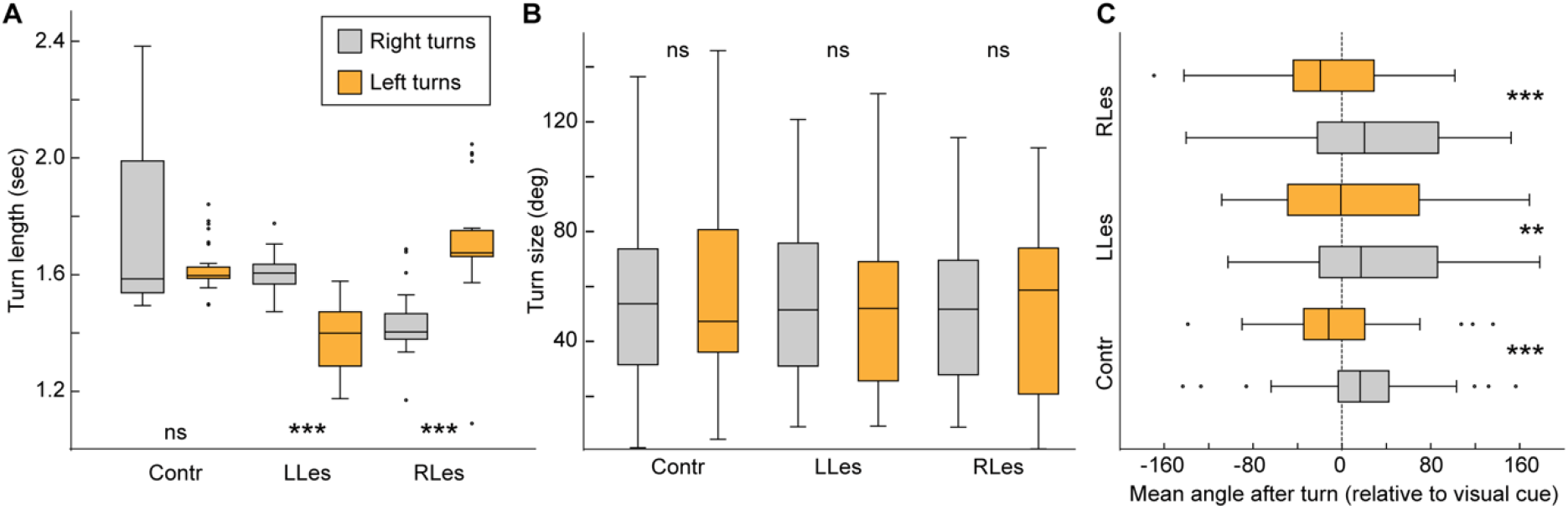
Lesions in the CX disrupt turning. **(A)** Control ants spent on average an equal amount of time turning to the left and right, whereas lesioned ants spent less time turning to the ipsilateral side of the lesion, i.e. right lesion ants spent less time turning to the right whereas left lesion ants spent less time turning to the left. Here and in the following panels, an average value from every ant is taken. **(B)** Size of turns did not differ between left and right turns in all three groups. **(C)** Angles at the end of the turns differed in all three groups between left and right turns, however, end points after both the left and right turns respectively did not differ between the three groups. Boxplots: median, 25th and 75th percentiles (edges of the boxes) and whiskers for extreme values not considered as outliers (circles). Wilcoxon tests: ns, not significant; **, p < 0.01; ***, p <0.001. For further statistics and sample sizes see Results.

## DISCUSSION

In recent years the CX has received attention as a brain area crucial for orientation in insects (Pfeiffer & Homberg 2014; Turner-Evans & Jayaraman 2016; Varga *et al.* 2017; Honkanen *et al.* 2019). The CX receives pre-processed sensory input that allows insects to orient themselves relative to external cues (Seelig & Jayaraman 2015). Furthermore, the CX can directly control movement (Bender *et al.* 2010). We show here that lesions in the CX lead to a lateralised disruption in ants’ turning behaviour (Figure 3) but that ants with lateralised CX lesions can still control their overall orientation relative to a visual cue (Figure 2). Below we consider two, non-mutually exclusive, impacts that our lesion might have had on the sensorimotor circuitry of the ants.

### Control of innate visual behaviour in the CX

Many insects show sensori-motor reflex behaviours, i.e. they perform particular motor behaviours in response to specific visual stimuli. Long vertical objects are attractive for many insects, including ants (fruit flies: (Wehner 1972; Gotz 1987; Strauss & Pichler 1998), locusts: (Wallace 1962), ladybirds: (Collett 1988), mantids: (Poteser & Kral 1995), ants: (Voss 1967; Heusser & Wehner 2002; Graham *et al.* 2003; Collett 2010; Buehlmann *et al.* 2020a; Buehlmann *et al.* 2020c; Buehlmann & Graham 2021)). Experiments in *Drosophila* have revealed that the CX is required for the flies’ innate visual responses (Bausenwein *et al.* 1994) and spatial learning paradigms in *Drosophila* revealed its importance for visual and spatial memory and directional decision making (Neuser *et al.* 2008; Ofstad *et al.* 2011; Kuntz *et al.* 2012; Seelig & Jayaraman 2013; Seelig & Jayaraman 2015). Neurons in the ellipsoid body map visual cues spatially (Seelig & Jayaraman 2013) and the lateralised ellipsoid body input in *Drosophila*, has been shown to be suitable for innate responses to bars (Dewar *et al.* 2017). In addition, ellipsoid body neurones have been shown to be important for short-term spatial memory, during which they encode the position of a target (e.g. an attractive visual cue) and can guide movement towards it (Neuser *et al.* 2008) even when temporarily invisible. Given the conservation of function in the CX across insects, it is likely that the ellipsoid body does play a similar role in ants. With lateralised lesions ants retained their innate attraction to the visual cue, albeit with lateralised disruption because the very wide field visual inputs from each visual hemisphere (Seelig & Jayaraman 2013; Dewar *et al.* 2017) are sufficient for overall path control, suggesting some redundancy in the visual inputs used for innate visual attraction.

### Control of turning behaviour by the CX

The CX is also involved in fine-scale control of steering including the control of walking speed (Bender *et al.* 2010; Martin *et al.* 2015) and turning (Guo & Ritzmann 2013). The symmetrical features of the CX along the medial-lateral axis might suggest a lateralized role of the CX (see e.g. (Pfeiffer & Homberg 2014). We show here that lesioned ants spent less time turning to the ipsilateral side of the lesion, i.e. right lesion ants spent less time turning to the right whereas left lesion ants spent less time turning to the left (Figure 3A). Ridgel and co-workers showed in cockroaches that off-centre CX lesions resulted in lateralised navigational deficits (Ridgel *et al.* 2007). More specifically, lesioned cockroaches lost the ability to turn in the ipsilateral direction relative to the lesion (Ridgel *et al.* 2007; Guo & Ritzmann 2013). Furthermore, CX stimulations experiments showed evoked turning to the ipsilateral side, i.e. stimulating the left CX evoked left turns and vice versa (Guo & Ritzmann 2013; Martin *et al.* 2015). Hence, the CX controls steering in a lateralized way, in both cockroaches and ants.

It has previously been suggested that visual control of orientation in wood ants involves ‘correction points’ that are synchronised with the underlying path ‘wiggle’ or sinuosity (Lent *et al.* 2010; Collett *et al.* 2014). This suggest that there may be some independence between the control of underlying path sinuosity (perhaps from the lateral accessory lobes; (Steinbeck *et al.* 2020)) and of visual corrections. Lateralised lesions in ants might not therefore knock-out the control of all turns in one direction but might diminish the visual control of those turns. However, redundancy in the visual inputs that allow for innate visual orientation, mean that ants’ overall paths are still directed to the visual cue.

## ACKNOWLEDGEMENTS

The research was funded by the Biotechnology and Biological Sciences Research Council (grant number: BB/R005036/1). We thank Yan Gu for help at the confocal microscope.

